# Predicting future regional tau accumulation in asymptomatic and early Alzheimer’s disease

**DOI:** 10.1101/2020.08.15.252601

**Authors:** Joseph Giorgio, William J Jagust, Suzanne Baker, Susan M. Landau, Peter Tino, Zoe Kourtzi, for the Alzheimer’s Disease Neuroimaging Initiative

## Abstract

The earliest stages of Alzheimer’s disease (AD) involve interactions between multiple pathophysiological processes. Although these processes are well studied, we still lack robust tools to predict individualised trajectories of disease progression. Here, we employ a robust and interpretable machine learning approach to combine multimodal biological data and predict future tau accumulation, translating predictive information from deep phenotyping cohorts at early stages of AD to cognitively normal individuals. In particular, we use machine learning to quantify interactions between key pathological markers (β-amyloid, medial temporal atrophy, tau and APOE 4) at early and asymptomatic stages of AD. We next derive a predictive index that stratifies individuals based on future pathological tau accumulation, highlighting two critical features for optimal clinical trial design. First, future tau accumulation provides a better outcome measure compared to changes in cognition. Second, stratification based on multimodal data compared to β-amyloid alone reduces the sample size required to detect a clinically meaningful change in tau accumulation. Further, we extend our machine learning approach to derive individualised trajectories of future pathological tau accumulation in early AD patients and accurately predict regional future rate of tau accumulation in an independent sample of cognitively unimpaired individuals. Our results propose a robust approach for fine scale stratification and prognostication with translation impact for clinical trial design at asymptomatic and early stages of AD.

**One Sentence Summary:** Our machine learning approach combines baseline multimodal data to make individualised predictions of future pathological tau accumulation at prodromal and asymptomatic stages of Alzheimer’s disease with high accuracy and regional specificity.

## Introduction

Alzheimer’s Disease (AD) is a protracted process with multiple pathophysiological events occurring well before clinical manifestations ^1^. Early phases involve molecular interactions between the β-amyloid (Aβ) and tau proteins. Quantifying these interactions is critical for establishing a mechanistic and precise account of the events that lead to progression of AD.

With the availability of PET imaging of Aβ and tau pathology in the brain, associations between AD related biomarkers can be examined in-vivo (for reviews: ^2–4^. These studies have shown that AD related cognitive impairment is dependent on cortical Aβ accumulation that facilitates the spread of tau into the neocortex resulting in neurodegeneration ^5^. In the absence of Aβ the presence of tau in the medial temporal lobe is insufficient to initiate widespread pathological neurodegeneration ^5, 6^. In addition, recent evidence linking the apolipoprotein E gene and tau suggests that the presence of the APOE4 allele worsens tau related neurodegeneration independent of Aβ ^7–9^. Given this model of the AD pathological cascade ^10, 11^, the staging of AD has shifted from a clinical syndromic diagnosis ^12, 13^ to a continuum of biomarker characteristics ^14^. In this framework, evidence of Aβ and pathological tau accumulation is sufficient to establish the diagnosis of AD. Cognitively unimpaired individuals (CN: cognitively normal) are classified as preclinical AD. Individuals with mild cognitive impairment (MCI) and these abnormal biomarkers are classified as having MCI due to AD or prodromal AD. These clinical syndromic definitions ^13^ have no discrete demarcations on cognitive scales, nor are they specific ^15, 16^ or sensitive ^17, 18^ to the underlying pathology of AD. As a result, there is a need for modelling approaches that predict longitudinal change in pathological biomarkers rather than simply a change in clinical labels based on syndromic diagnosis.

Here, we employ machine learning to quantify the multivariate relationships between key pathological markers that underly the pathogenesis of AD: APOE 4 genotype, Aβ, tau and neurodegeneration. We have developed a machine learning approach that derives a single prognostic index from APOE 4 genotype with measures of Aβ derived from either [^18^F]-florbetapir (FBP) or [^11^C]-PiB PET scans and a continuous measure of medial temporal atrophy derived from structural MRI ^19^. Using this approach, we previously demonstrated that the model derived prognostic index is predictive of individualised rates of future memory change in MCI patients ^19^. Here, we use our machine learning approach to test whether a prognostic index derived from baseline data classifies and stages early AD (i.e. CN and MCI) individuals based on future pathological tau accumulation.

Recent converging evidence highlights that patterns of tau spread (measured in-vivo by longitudinal FTP-PET) are robust across early AD cohorts ^20–22^. This stereotypical spreading pattern for early AD (i.e. Aβ positive individuals who are cognitively unimpaired or individuals with intermediate levels of baseline tau) shows that tau initially accumulates within the temporal cortex then spreads to the superior and medial regions of the parietal cortex prior to severe cognitive impairment ^20–22^. Here, we employ our machine learning approach to predict the future rate of tau deposition in two independent samples and examine the clinical utility of our approach in the context of clinical trials.

## Results

### Machine learning derives an accurate prognostic index from multimodal biomarkers

We used a trajectory modelling approach based on the Generalised Matrix Learning Vector Quantisation (GMLVQ) machine learning framework (GMLVQ-scalar projection ^19^) to generate a prognostic index as a single numerical descriptor (scalar projection) from three biological markers measured at baseline: cortical Aβ measured using PET, medial temporal grey matter density measured using T1 weighted MRI and APOE 4 genotype. This approach derives a continuous prognostic metric by training a learning model with classes determined by longitudinal diagnostic labels.

In particular, we trained the model on baseline (defined as the date of Aβ PET scan) data from the Alzheimer’s Disease Neuroimaging Initiative (ADNI) 2/GO cohort. We determined two classes for training the algorithm: a) Stable Condition (SC, n=145): cognitively normal individuals who remain stable for 4+ years following baseline, b) Early AD (EAD, n=162): individuals with unstable diagnosis (i.e. those with MCI or CN diagnosis at baseline who are subsequently diagnosed with dementia or those who had reverted from a diagnosis of dementia prior to baseline to MCI at baseline). We included individuals who reverted in the EAD group because they are presumably at an earlier stage of AD than patients with a stable diagnosis of dementia. An additional 181 participants with stable diagnoses of dementia were included for comparison. Comparing the unimodal distributions of the SC and EAD groups shows that the EAD group has greater AD pathology than the SC group across the three biological predictors (Aβ: t(305)=14.04; p<0.0001, medial temporal grey matter density t(305)=-11.02; p<0.0001, APOE 4 χ2(1,305)=39.02, p<0.0001). Contrasting medial temporal grey matter density for the EAD group with a group of individuals with a stable diagnosis of dementia, we showed that the EAD group has greater medial temporal grey matter density than a demented group (t(341)=-18.15, p<0.0001), suggesting that the EAD group is at an earlier pathophysiological stage (**Figure 1**).

**Figure 1:**
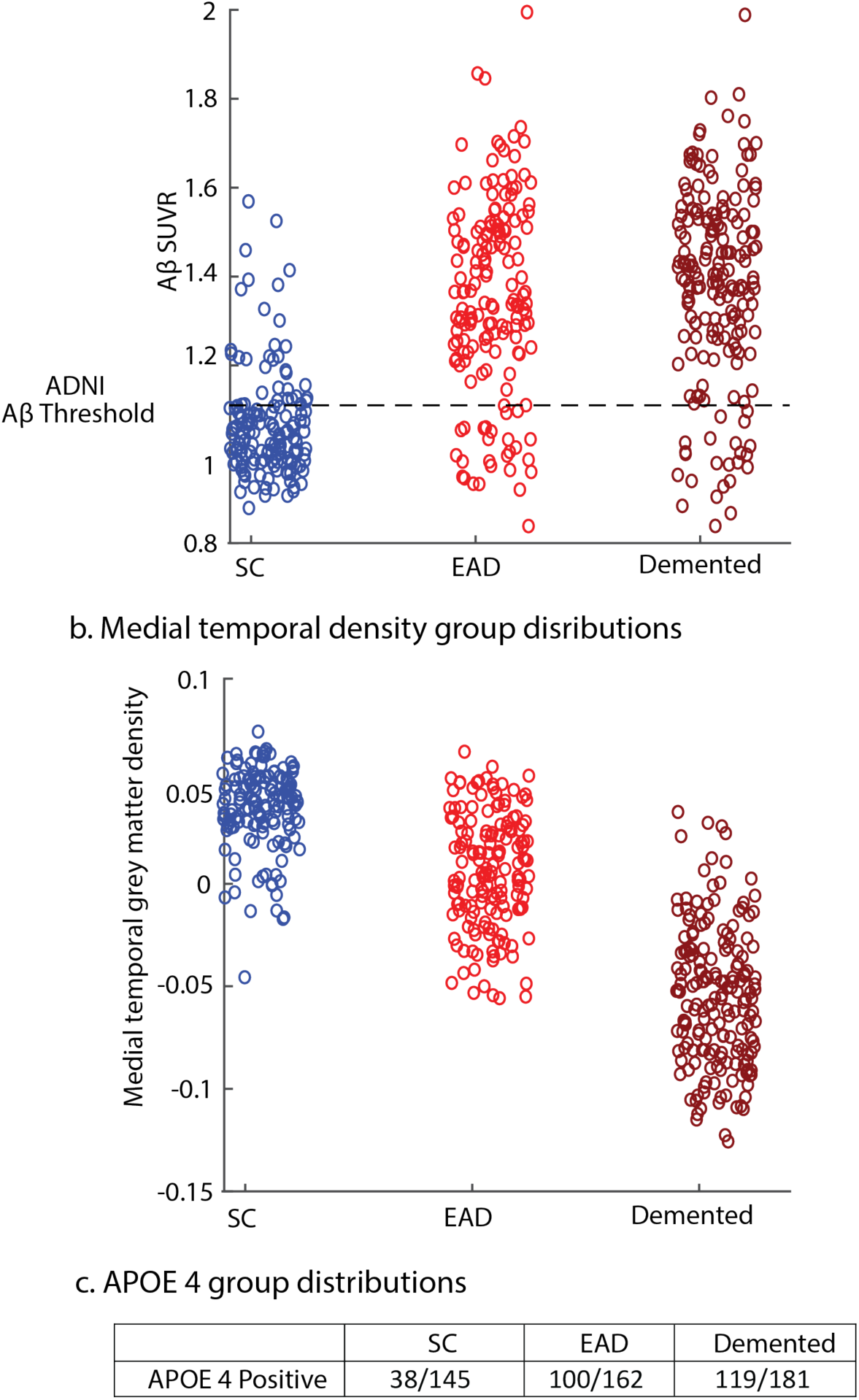
Distributions of biological predictors for SC, EAD and Demented groups. distribution of the biological predictors used in the GMLVQ-scalar projection model for the ADNI2/GO sample. Blue dots indicate individuals in the Stable Condition group, red dots indicate individuals in the Early AD group and maroon dots indicate individuals in the Demented group. **a.** distribution of FBP (Aβ) SUVR values, the dashed horizontal line indicates the ADNI threshold of Aβ positivity (1.11). **b.** distribution of medial temporal grey matter density. **c** proportions of APOE 4 positive individuals in the Stable Condition, Early AD and Demented groups.

We trained our GMLVQ-scalar projection model to learn the multivariate relationship between the three baseline biological predictors (metric tensor) and the location in multidimensional space (prototype) that best discriminates between stable condition and early AD individuals. We then determined the distance of each individual from the stable condition prototype along the axis that is predictive of future diagnosis (i.e. prognostic axis from stable towards EAD). That is, we derived a scalar value from the projection of any sample point along the prognostic axis.

Using random resampling to split the ADNI2/GO sample into training and test sets we demonstrated that the model classifies SC vs. EAD individuals with cross-validated class-balanced accuracy of 86.4%. Further, using logistic regression we showed that an individual is more than 50% likely to be classified as EAD if they have a scalar projection greater than 0.4. Comparing the scalar projection to the three biological markers showed that the multimodal scalar projection captures predictive variance in each of the unimodal predictors (scalar projection vs. Aβ: R^2^=77%, p<0.0001; scalar projection vs. atrophy: R^2^=37%, p<0.0001; scalar projection (APOE 4 - /+): t(305)15.4, p<0.0001) (**Figure 2**).

**Figure 2:**
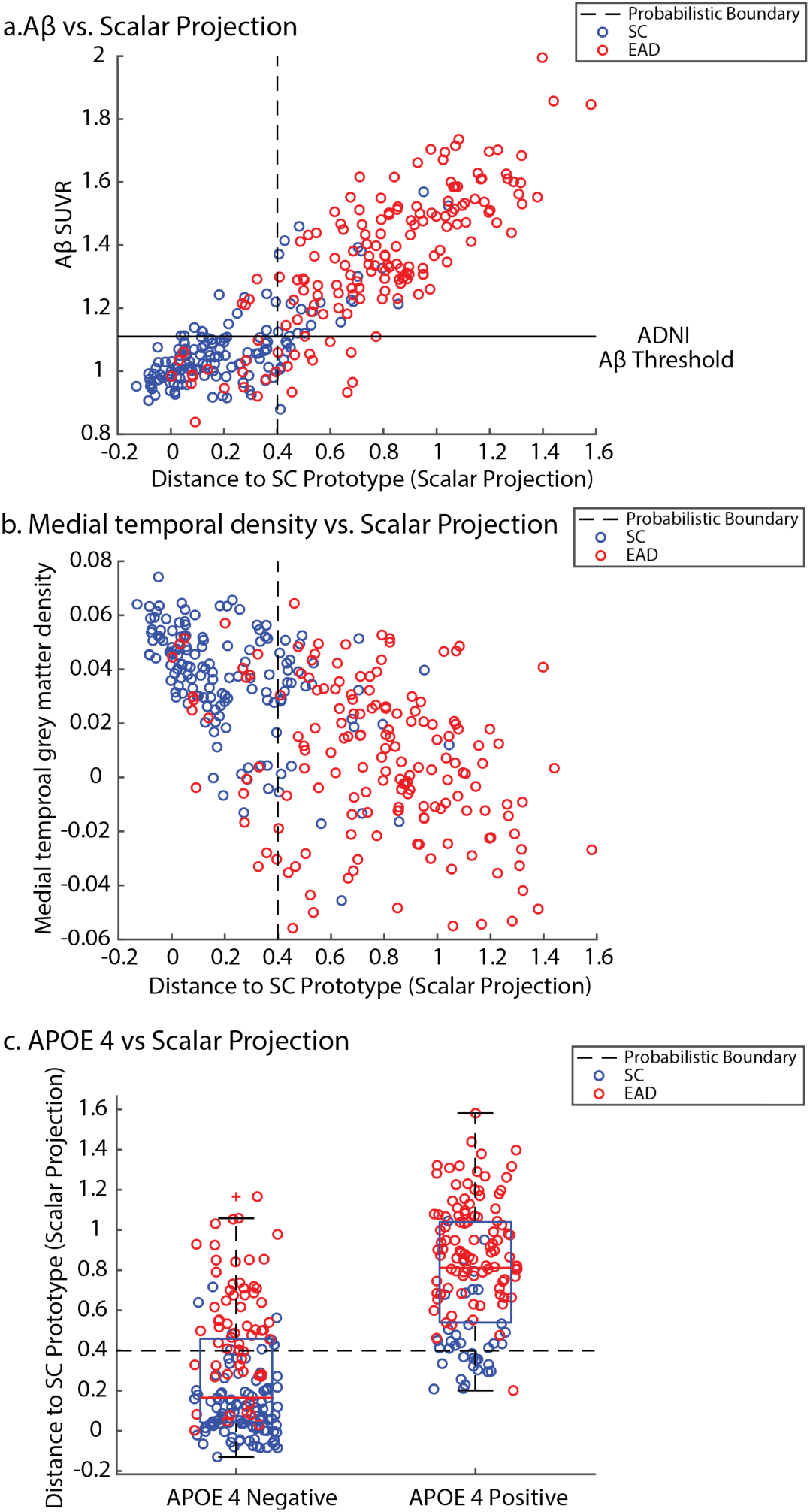
Relationship of scalar projection with biological predictors. relationship of scalar projection with the biological predictors for the ADNI2/GO sample. Blue dots indicate individuals in the Stable Condition (SC), red dots indicate individuals in the Early AD (EAD) group. **a.** Relationship of scalar projection with FBP SUVR (Aβ), the solid horizontal line indicates the ADNI threshold for Aβ positivity (1.11), the dashed vertical line indicates the learnt probabilistic boundary that separates SC from EAD (0.4). **b.** relationship of scalar projection with medial temporal grey matter density, the dashed vertical line indicates the learnt probabilistic boundary that separates SC from EAD (0.4). **c.** relationship of scalar projection with APOE 4 status, the dashed horizontal line indicates the learnt probabilistic boundary that separates SC from EAD (0.4).

Next, we used the model trained on ADNI2/GO data to derive the scalar projection for individuals from two independent cohorts: a) ADNI 3 (CN=72, MCI=43) b) Berkeley Aging Cohort Study (BACS) (CN=56). The scalar projection classified 61 ADNI3 participants and 33 BACS participants as SC, with 54 ADNI3 and 23 BACS participants classified as EAD. (i.e. scalar projection greater than 0.4) (**Figure 3**). This scalar projection was weakly related to baseline age in ADNI 3 (r(113)=0.28, p=0.003) but not in BACS (r(54)=0.187, p=0.168) and it was unrelated to education (BACS: r(54)=-0.011, p=0.94, ADNI 3: r(113)=-0.014, p=0.88) or sex (BACS: t(54)=-1.52, p=0.136, ADNI 3: t(113)=-0.582, p=0.56). Finally, for the ADNI 3 sample we compared whether a classification performed using the multimodal scalar projection is similar to syndromic clinical diagnosis. Comparing the classification of SC vs. CN, and EAD vs. MCI (**Figure 3b**) showed that agreement was not significantly greater than chance (Cohens kappa κ=0.17 [-0.0126, 0.3524] p=0.084). That is, the clinician-based diagnosis and the multimodal scalar projection have poor agreement for differentiation of SC and EAD.

**Figure 3:**
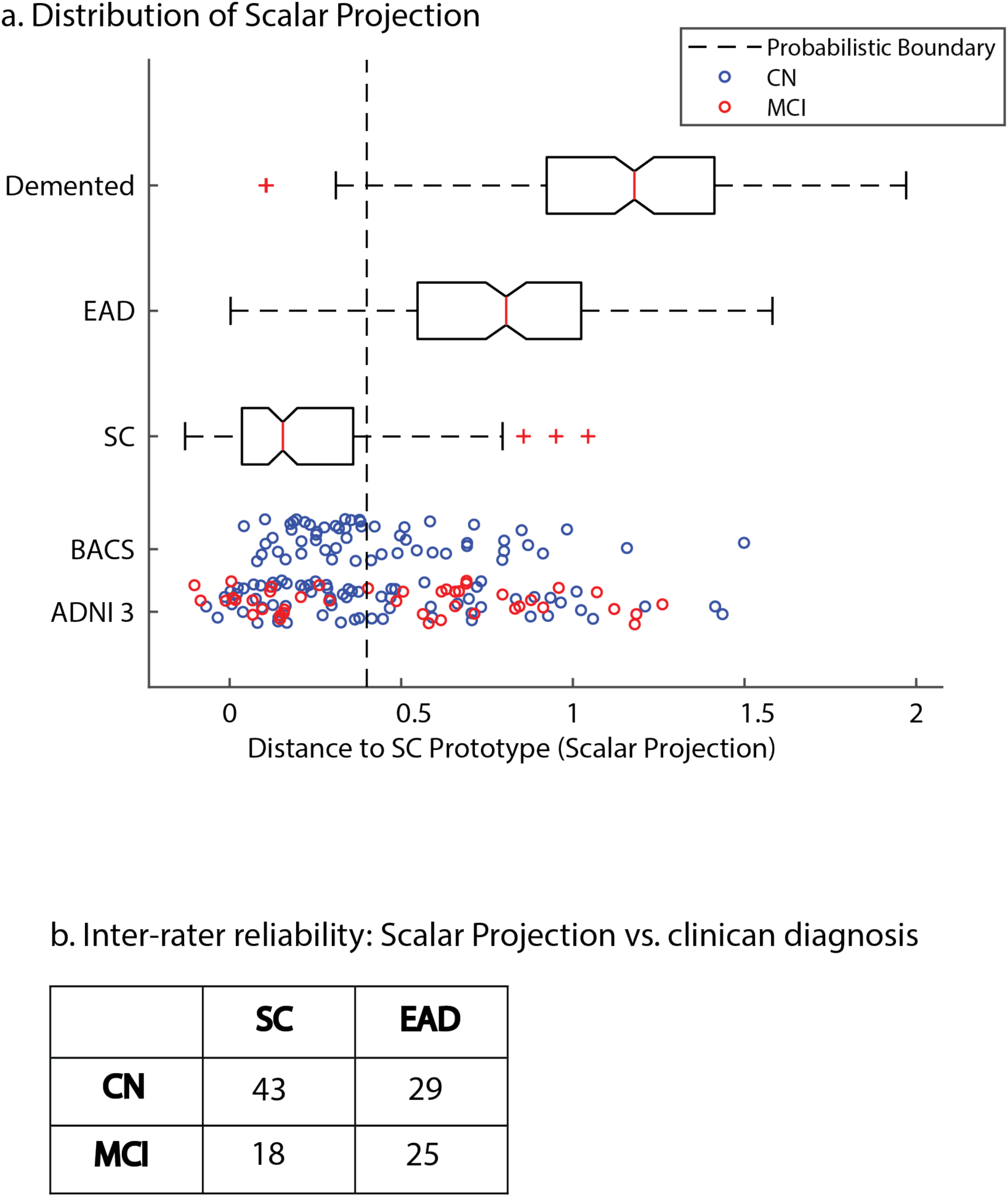
Distribution of the scalar projection for the ADNI2/GO, ADNI3 and BACS samples. a. The distribution of the scalar projection for individuals from ADNI2/GO in the SC (Stable Condition), EAD (Early AD) and Demented. Blue dots indicate individuals from ADNI 3 / BACS who are cognitively normal (CN) at baseline. Red dots indicate individuals from ADNI 3 who have Mild Cognitive Impairment (MCI) at baseline. The dashed vertical black line represents the probabilistic boundary used to classify SC vs EAD (0.4), all individuals to the right of the line are classified as EAD. **b.** inter-rater reliability when diagnosing early AD based on biological predictors (i.e scalar projection) or a clinical diagnosis based on syndromic definitions (i.e. CN or MCI).

### Individuals classified as early AD accumulate tau more rapidly that stable individuals

We used longitudinal FTP-PET to compare regional longitudinal tau accumulation for individuals classified as SC vs. EAD from the ADNI 3 sample. We extracted regional SUVR values from 36 Desikan-Kilany ROIs and calculated annualised rate of change per ROI for each individual. ROIs were grouped together in order to approximate the topographical distribution of tau in the Braak Staging scheme, as previously described ^23^. For individuals with 2 FTP-PET scans we used the difference between the follow-up and baseline FTP-PET scans divided by the time interval in years from baseline. For individuals with 3 or more FTP-PET scans we used the linear least squares fit of time from baseline vs. ROI SUVR.

We averaged the annualised rate of tau accumulation within each of the 36 Desikan-Kilany ROIs for SC and EAD groups then contrasted the global rate of tau accumulation for SC vs EAD (i.e. independent samples t-test across ROIs for SC vs EAD).We observed a medium to large effect when comparing global tau accumulation for SC vs EAD (t(70)=2.96, p=0.004, Cohens d=0.7), with the EAD group accumulating global cortical tau 2.8 times faster than SC (**Supplementary Table 1**). Further, testing which regions significantly accumulated tau (i.e. rate of accumulation significantly greater than 0; one sample (i.e. SC or EAD) one tail t-tests within each ROI) we showed that EAD individuals accumulate tau primarily in Braak stages 4 and 5 ROIs (**Supplementary Table 1**, **Figure 4**). In contrast, individuals classified as SC did not show clear future tau accumulation across cortical regions, with a marginal effect in the middle temporal ROI t(60) =1.71, p=0.047) (**Figure 4bii)**. There was no difference in cognitive change (as measured by future annualised change in PACC) over the same time period between individuals classified as SC(mean=-0.161/year) vs. EAD(mean=-0.6/year) (t(100)=-1.04, p=0.30). However, individuals classified as EAD showed significant worsening (i.e. rate of PACC change significantly less than 0) in future cognitive ability (one tail t-test t(47)=-1.87, p=0.034).

**Figure 4.**
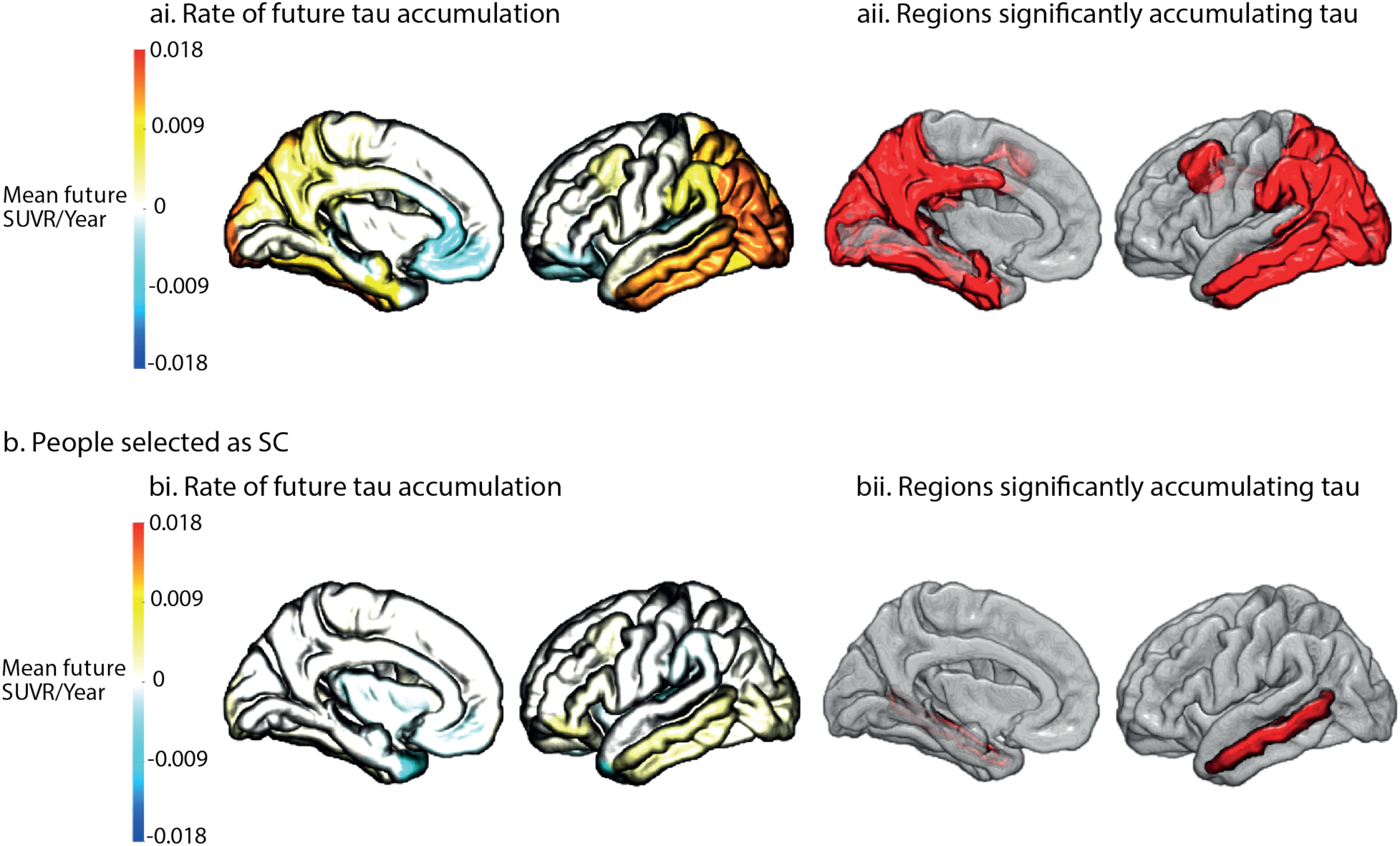
Regional future rate of tau accumulation. The future annualised rate of tau accumulation for ADNI 3 individuals across the 36 Desikan Kilany ROIs. ai. Mean future annualised rate of tau accumulation for individuals classified as the Early AD (EAD). aii The regions in red are significantly (p<0.05 uncorrected) accumulating tau for individuals classified as Early AD (EAD). bi. Mean future annualised rate of tau accumulation for individuals classified as Stable Condition (SC). bii The regions in red are significantly (p<0.05 uncorrected) accumulating tau for individuals classified as Stable Condition (SC).

Next, we compared how sensitive a syndromic classification of CN vs MCI is to future changes in tau accumulation. Averaging the annualised rate of tau accumulation within each of the 36 Desikan-Kilany ROIs for CN and MCI groups we contrasted the global rate of tau accumulation for CN vs MCI groups (i.e. independent samples t-test across ROIs for CN vs MCI). We observed a small to medium effect when comparing global tau accumulation between CN and MCI groups (t(70)=2, p=0.05, Cohens d=0.48), with MCI individuals accumulating global cortical tau 1.9 times faster than CN individuals. Further, testing which regions significantly accumulated tau (i.e. rate of accumulation significantly greater than 0; one sample (i.e. CN or MCI) one tail t-tests within each ROI) we showed that both CN and MCI individuals are significantly accumulating tau, with a high degree of overlap across AD susceptible regions in the temporal and posteromedial cortices (**Supplementary Table 2**, **Supplementary Figure 1**). Taken together this highlights that a stratification based on syndromic diagnosis has poorer sensitivity and specificity to future tau accumulation than the biological classification of SC vs EAD based on the multimodal scalar projection.

### Fewer patients are required to detect meaningful change in tau accumulation than cognition

To examine the clinical utility of regional longitudinal tau accumulation as an outcome measure, we contrasted the sample size required to observe cognitive decline vs. tau accumulation for individuals classified as EAD based on the scalar projection. We defined a clinically meaningful change as a 25% reduction in rate of change of either regional tau accumulation or PACC change. For individuals classified as EAD we calculated that the required sample size (for a significance level of p=0.05 at a power of a=0.8) to detect a 25% reduction in rate of PACC change is n=1730. However, to detect a 25% reduction in regional future tau accumulation in the selected areas (**Figure 4c**) at the same power, an average sample size of n=1127 is required. Thus, using future rate of tau accumulation as a clinical outcome measure instead of future rate of cognitive decline delivers a reduction in required sample size of 35%.

### Using multimodal biomarkers reduces sample size to detect a meaningful change in tau accumulation

Next we compared prediction of longitudinal tau accumulation for individuals classified as EAD (n=54) based on the scalar projection vs. those classified based on Aβ (n=61 Aβ positive). We observed that individuals classified as EAD accumulated global cortical tau (i.e. mean rate of tau accumulation across the 36 Desikan-Kilany ROIs) 1.5 times faster than individuals who were defined only as Aβ positive (**Supplementary Table 2)**.

We next tested for the sample size needed to detect a 25% decrease in rate of future tau accumulation (given significance level of p=0.05 at power of a=0.8) for EAD vs. Aβ positive individuals within regions shown to significantly accumulate tau for individuals classified as EAD (**Figure 4c**). We observed on average a 47% reduction in sample size when stratifying based on EAD classification (n=1127) vs. Aβ positive alone (n=2146). We repeated these calculations using regions in which Aβ positive individuals were shown to significantly accumulate tau **(Supplementary Table 3)**. This showed an average of 30% reduction in sample size when stratifying based on EAD classification (n=831) vs. Aβ positive alone (n=1190) (**Figure 5**). These results provide evidence for the clinical utility of stratification based on the EAD classification using the scalar projection derived from multimodal data compared to Aβ status alone.

**Figure 5.**
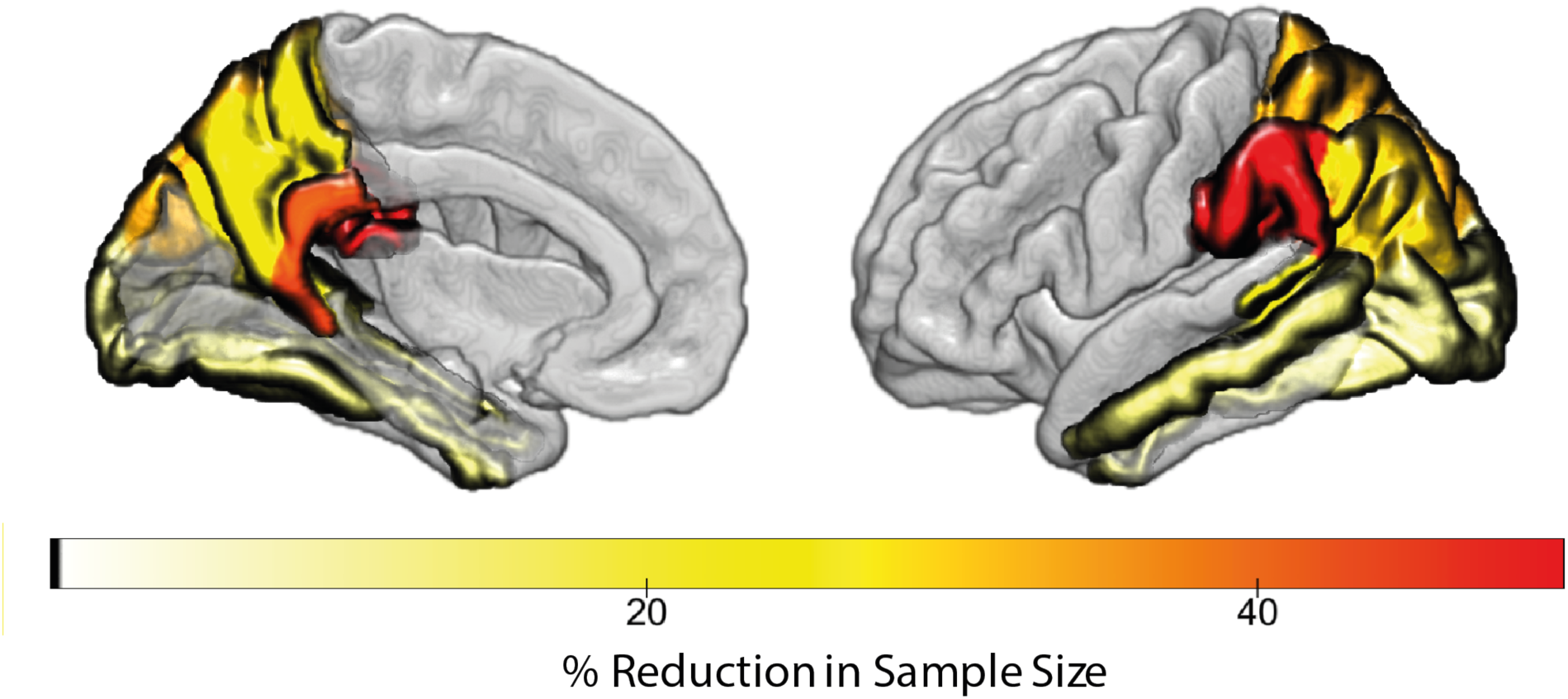
Reduction in sample size to observe clinically meaningful change when stratifying using multimodal data vs. Aβ positive only. percentage reduction of sample size required to observe a 25% reduction in tau accumulation per region for EAD classification based on multimodal data (EAD) vs. Aβ status alone. For example, to observe a 25% reduction in tau accumulation in the middle temporal cortex 14% fewer individuals are required when stratifying based on EAD classification vs Aβ status alone. In medial and superior parietal regions (e.g. Precuneus or superior parietal) a sample size reduction of 26% and 33% is achieved when stratifying based on EAD vs Aβ status alone.

### Scalar projection predicts individual variability in trajectories of regional tau accumulation

Using the ADNI 3 sample we fit linear regression equations to test whether the scalar projection (derived from the model trained on ADNI2/GO) predicts individual variability in future tau accumulation (**Supplementary Table 4)**. Of the 15 regions that were shown to significantly accumulate tau (**Figure 4aii)** 13 showed a significant relationship of the scalar projection and individual rates of future tau accumulation (**Figure 6a**), explaining up to 30% of variance in the temporal cortex and 20% in superior and medial regions of the posterior parietal cortex (**Figure 6b**).Further, we generate two aggregate regions by averaging future rate of tau accumulation across a.) temporal, and b.) parietal ROIs significantly accumulating tau. Within these aggregate regions we observed a significant relationship between the scalar projection and future tau accumulation describing 18% of the variance in the temporal aggregate region and 27% in the parietal aggregate region (**Figure 6c**).

**Figure 6.**
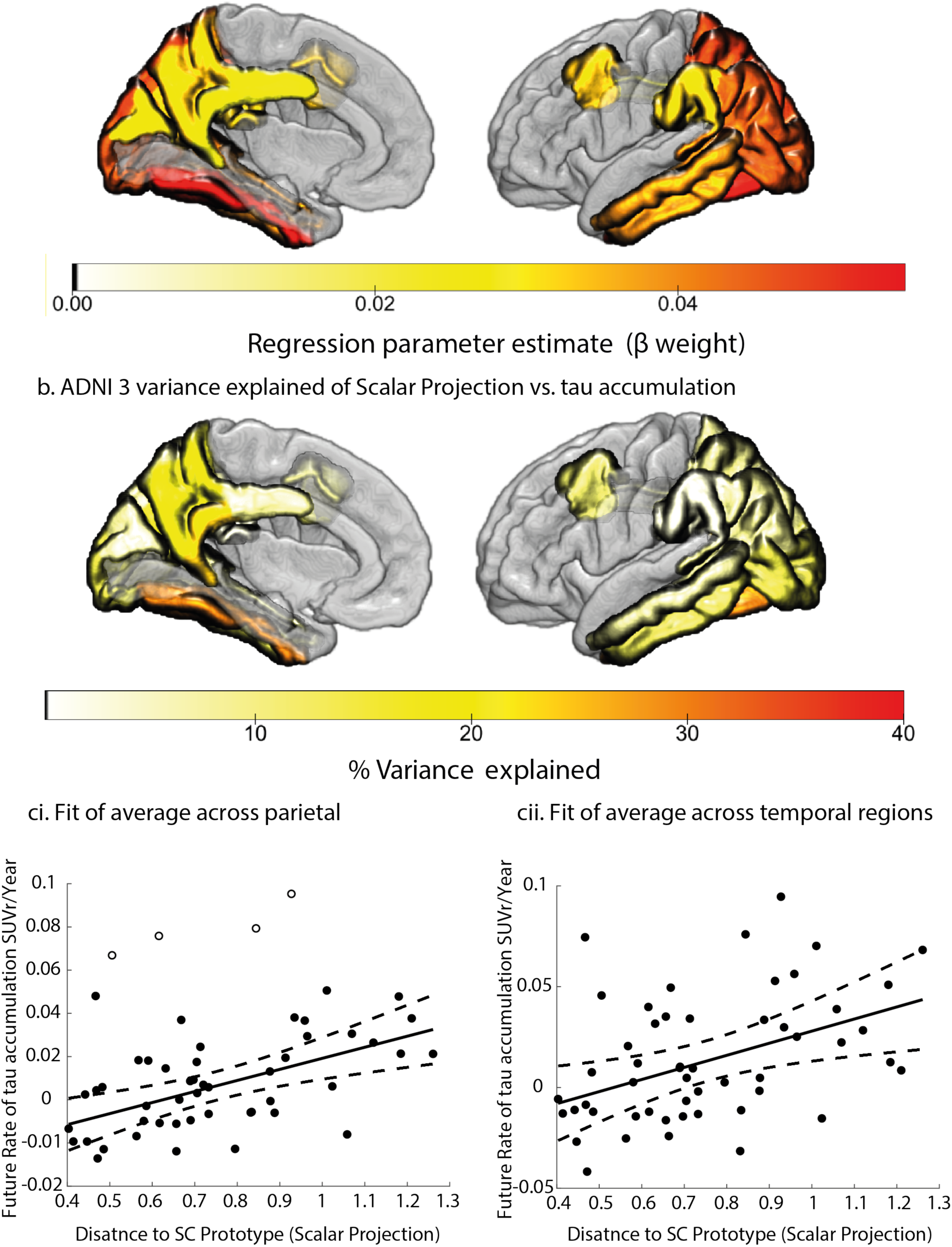
Predicting individual variability in future tau accumulation in ADNI 3 early AD a. Significant (p<0.05 uncorrected) regional parameter estimates from linear regressions to predict future rate of tau accumulation for individuals classified as Early AD (EAD). The colour scale indicates the parameter estimate for the slope of the regression fit of the scalar projection with regional future tau accumulation. **b.** percentage of variance explained when using the scalar projection to predict future rate of tau accumulation for individuals classified as EAD from the ADNI 3 cohort. **c** regression fits of the scalar projection with future rate of tau accumulation for; **i)** an aggregate parietal region comprised of; supramarginal gyrus, inferior parietal lobe, superior parietal lobe and precuneus (R^2^=27%), **ii)** an aggregate temporal region comprised of; fusiform gyrus, middle temporal gyrus, inferior temporal gyrus and banks of the superior temporal sulcus (R^2^=18%). The central black line indicates the regression line for the regression fit; the dashed lines indicate the 95% confidence intervals for this regression line. Outliers identified by the Robust Correlation toolbox are shown as unfilled black dots. Two outliers based on the scalar projection is not shown for illustrative purposes.

### Individual trajectories of pathological tau accumulation are accurately predicted in an independent asymptomatic cohort

To test the robustness of these regional fits we generated individualised predictions of future tau accumulation for CN individuals classified as EAD from the BACS sample. Using the model trained on ADNI2/GO individuals, we derived the scalar projection from baseline multimodal biological data in the BACS sample (**Figure 3**). To predict the future rate of tau accumulation in this sample, we used the linear regression models relating the scalar projection to rate of future tau accumulation derived from the 13 regions that showed significant fits in the ADNI 3 sample (**Figure 6a**, **Supplementary Table 4**). Comparing the predicted and real future rate of tau accumulation for BACS EAD individuals we observed that 7 ROIs (**Figure 7b**) pass the critical value of variance explained (R^2^>17%) for significant correlations with a comparable sample size (n=23). These individualised predictions of future tau accumulation explain up to 39% of the variance in the temporal cortex and 32% in superior and medial regions of the posterior parietal cortex (**Supplementary Table 5**, **Figure 7**). Further, using the regression models within the temporal and parietal aggregate regions (**Figure 6c**) for EAD individuals the predicted future tau accumulation explains 30% of the variance in temporal, and 20% in the parietal regions of observed future rate of tau accumulation (**Figure 7c**).

**Figure 7.**
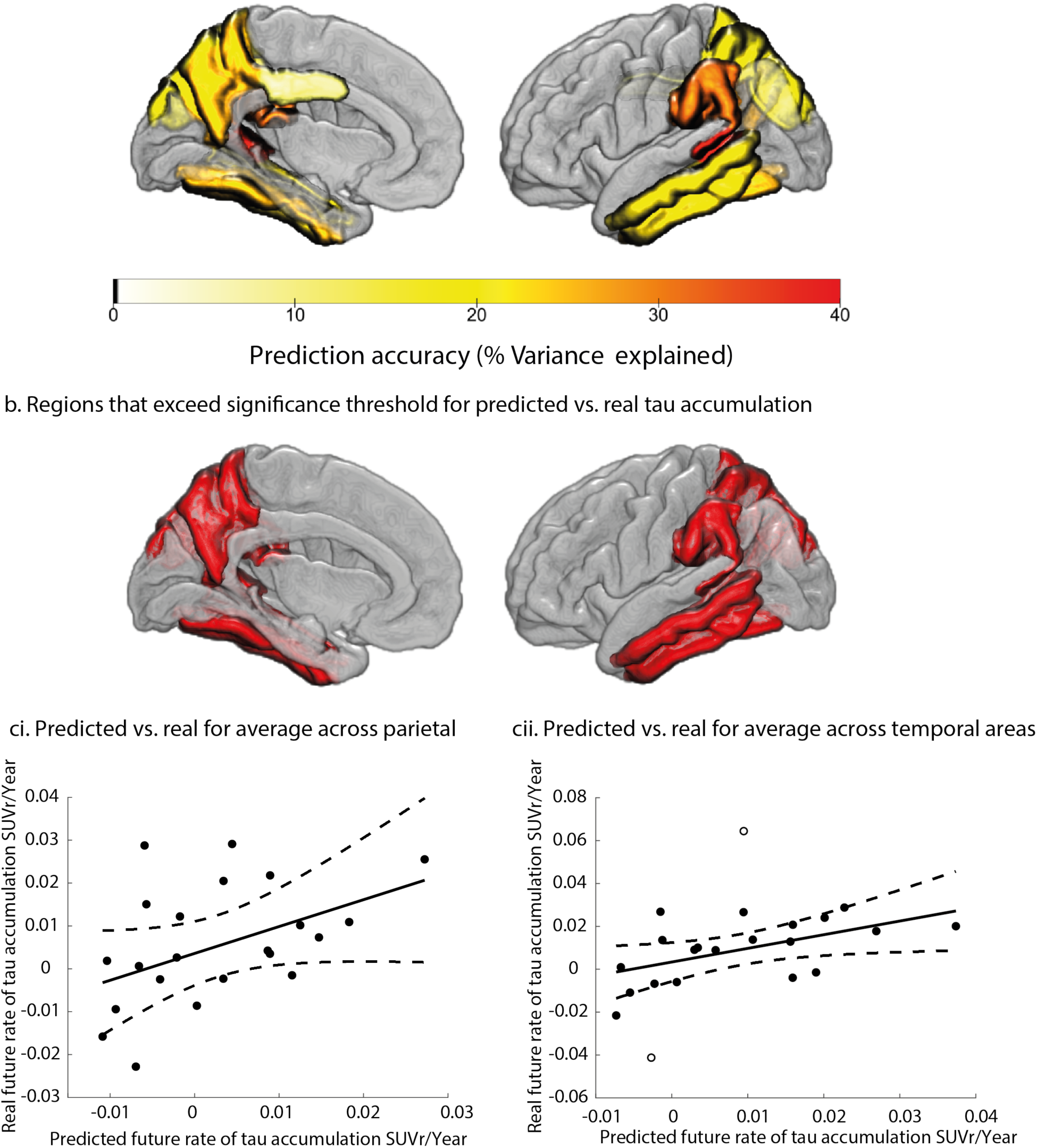
Prediction accuracy of future tau accumulation in BACS early AD. Comparing predicted tau accumulation to observed tau accumulation for individuals classified as Early AD (EAD)in the BACS cohort. **a.** percentage of shared variance of the predicted regional future tau accumulation and the observed future tau accumulation for EAD individuals within each of the selected ROIs (Figure 6a.). **b.** regions with shared variance of predicted future tau accumulation and observed tau accumulation that exceeds the critical value for significance of p<0.05 (i.e. R^2^(21)>17%) **ci.** Fit of the predicted future rate of tau accumulation to the observed tau accumulation within the aggregate regions shown in (Figure 6c). accuracy of predicted future tau accumulation within the parietal aggregate region (R^2^=20%) **cii**. accuracy of predicted future tau accumulation within the temporal aggregate region (R^2^=30%). The central black line indicates the regression line for the regression fit; the dashed lines indicate the 95% confidence intervals for this regression line. Outliers identified by the Robust Correlation toolbox are shown as unfilled black dots. One outlier based on the scalar projection is not shown for illustrative purposes.

Finally, we showed that for individuals who were classified as SC from the BACS sample no accurate prediction can be made with the predicted regional future tau accumulation describing on average only 1.1% of the observed variance in future tau accumulation **(Supplementary Table 5**). Therefore, our predictions are robust and specific for staging individuals who transition from non-pathological aging to Alzheimer’s Disease.

## Discussion

Here, we employ a robust and transparent machine learning approach to combine continuous information across AD biomarkers to predict pathological changes in tau accumulation in early and asymptomatic stages of AD. We use well characterised AD biomarkers (Aβ, medial temporal grey matter density, APOE 4) to generate a simple and straightforward numerical prognostic index (scalar projection) for stratification. Using this multimodal index derived from baseline data, we predict future tau accumulation, a known pathological driver of AD progression. We demonstrate that this multimodal prognostic index of future tau accumulation is a more sensitive tool for patient stratification than Aβ status alone. Further, our prognostic index predicts individualised spatially specific changes in tau accumulation in an out-of-sample group of cognitively normal individuals, enabling fine stratification and staging at asymptomatic stages (i.e. before clinical symptom occurrence). This strong predictive ability has important implications for both understanding the mechanisms of AD progression and developing methods for clinical trials.

In particular, employing our recently developed machine learning approach ^19^, we derived a prognostic index (scalar projection) from biomarkers measured at baseline that accurately classified individuals in the early stages of AD. Using this model derived prognostic index we showed that individuals classified as early AD will accumulate tau in a topography stereotypical of early AD ^20–22^. Consistent with previous work, we successfully stratified individuals based on the pattern of future tau accumulation in early AD, accurately reproducing the topography reported in numerous independent cohorts corresponding to the proposed “meta-ROI” for tau quantitation ^20–22^. Extending beyond previous work, we show that our multimodal prognostic index is more sensitive for predicting tau accumulation compared to a unimodal measure (i.e. Aβ positive alone). Further, we demonstrate that individuals who are classified as early AD using this index accumulate tau at 1.5 times the rate of Aβ positive individuals. Thus, an optimal combination of a small number of variables at baseline can select those at greatest risk of rapid tau accumulation.

Our approach has strong clinical relevance in two main respects. First, we show that using the rate of tau accumulation to classify early AD individuals results in 35% reduction in the sample size necessary for detecting a clinically meaningful change relative to the gold standard cognitive instrument (PACC ^24^). This is consistent with previous work showing that a smaller sample size is required to detect clinically meaningful change in tau accumulation within the “meta-ROI” for tau accumulation than using a cognitive endpoint ^22^. In addition, however, using the multimodal prognostic index vs. Aβ status alone to stratify individuals reduces the sample size required to observe a clinically meaningful change in the stereotypical pattern of pathological tau accumulation by 46%. The benefit of combining multimodal data for stratification in early AD has been previously reported in the context of future changes in cognition ^25–30^. These previous studies have shown that grey matter atrophy and cortical Aβ burden relate to separable patterns of future cognitive decline ^25, 26, 29, 30^. Given the known association between longitudinal changes in tau and cognitive decline in preclinical AD ^6^, our results further support the benefit of combining continuous values of Aβ and medial temporal grey matter density for prognostication in early AD. Thus, our multimodal prognostic index has strong potential to benefit clinical trial design targeting the early and asymptomatic stages of AD.

Extending beyond binary classifications, we show that our prognostic index predicts individual variability in future regional tau accumulation within regions that are known to be affected in early AD. We show that these individualised predictions generalise to an independent sample of cognitively normal community dwelling individuals. Further, we show within this asymptomatic population that these predictions are specific to individuals who are classified as early AD. Previous studies have investigated whether antecedent changes in Aβ relate to cross sectional tau deposition ^31, 32^, our work differs from these previous approaches as we modelled cross sectional interactions to predict future changes in tau. When investigating individual variability in future tau accumulation, previous work has focussed primarily on associations between simultaneous changes in pathophysiology ^6, 33, 34^. In contrast, our multimodal modelling approach allow us to make explicit individualised predictions specific to early AD, translating predictions from a research cohort to an asymptomatic community sample.

We consider the following potential limitations of our machine learning modelling approach. First, our model is trained on grey matter atrophy in the medial temporal lobe. Previously we established that this grey matter value relates to memory deficits, baseline tau and separates individuals who are stable MCI from progressive MCI ^19^. However, it is likely that we captured information specific to typical amnestic AD populations, as we derived this measure of atrophy based only on ADNI data. As dissociable patterns of tau spreading have been observed for atypical AD variants (i.e. posterior cortical atrophy and logopenic progressive aphasia) ^35^ additional measures of atrophy might be necessary for a larger scale predictive model. Second, we did not extensively investigate the predictive power of different markers. Neuropsychological data are shown to be predictive of MCI progression to dementia due to AD ^19, 36–41^, with predictive performance improving when incorporating biological information ^19, 42–44^. Further, recent studies have shown that blood based AD biomarkers have substantial predictive power in modelling AD trajectories ^45, 46^. Given the expense of in-depth neuroimaging phenotyping and the low frequency of community based cognitively normal individuals with appropriate AD biomarkers, a model that integrates cheap and readily available predictors (i.e. cognitive and blood plasma) is optimal for stratification in the earliest stages of AD. Finally, in this work we investigated the predictive performance of a single machine learning approach (GMLVQ-scalar projection). Machine learning in AD research is a relatively new and rapidly expanding field with most research investigating binary changes in diagnosis from baseline (for review: ^47–51)^. Direct comparison of diverse approaches remains challenging as cross validation methodology, sample sizes and sample heterogeneity have a significant effect on model performance metrics (i.e. accuracy or receiver operator characteristics)^52^. Although comparison of multiple machine learning approaches built on the same data may not be valid (see no free lunch theorem ^53^), prediction challenges offer a more unbiased approach for determining the efficacy of prediction models (e.g. TADPOLE ^54^). We validated our model trained on research cohort data by testing predictions in an independent cognitively normal sample; further testing in large cohorts and clinical data will increase our model generalisability.

Despite these potential limitations, we demonstrate that our multimodal modelling approach is more sensitive in capturing AD related pathology (i.e. EAD) than a classification based on syndromic labels. This poor sensitivity and specificity of syndromic labels to AD pathology ^15–18^ led to the introduction of a biological framework for AD classification ^14^. Using our machine learning approach we capitalise on longitudinal data with the clinical categorisation of participants following the 2011 NIA-AA syndromic definitions of AD ^12, 13^ (e.g ADNI ^55^) to make sensitive and specific classifications of early AD based on pathophysiology. Further, our approach combines continuous biological measures to capture trajectories for individuals that may be on the threshold of biomarker positivity but who are likely to follow AD related trajectories ^56^. Taken together our approach is well suited for harmonising longitudinal data collected using diagnostic criteria by combining continuous biomarkers into a biologically informative prognostic index in an interpretable and clinically meaningful way.

In sum, employing a transparent machine learning approach to derive a simple numerical scalar value from baseline measurements (Aβ, brain atrophy and APOE genotype) has strong predictive value for the subsequent topography and rate of pathological tau deposition. As tau accumulation represents a key disease stage that mediates effects of AD pathology on cognition, our findings link these baseline data to fundamental mechanisms of AD pathogenesis. Further, individualised prediction of future pathological tau accumulation has strong clinical relevance for prognosis and patient selection for inclusion in clinical trials. Thus, our findings demonstrate strong potential for machine learning approaches to capitalise on rich multimodal data, reducing their complexity and delivering tools with high predictive ability and clinical utility.

## Materials and Methods

### Study Design and Participants

Three separate cohorts were used to generate and test predictive models of regional future tau accumulation.

Two cohorts were drawn from the ADNI database: ADNI2/GO and ADNI 3 (adni.loni.usc.edu). ADNI was launched in 2003 as a public-private partnership, led by Principal Investigator Michael Weiner, MD. A major goal of ADNI has been to examine biomarkers including serial magnetic resonance imaging (MRI), and positron emission tomography (PET), with clinical and neuropsychological assessment to predict outcomes in mild cognitive impairment (MCI) and early Alzheimer’s disease (AD).

A third validation cohort was taken from the Berkeley Aging Cohort Study (BACS). This cohort is comprised of a convenience sample of community-dwelling cognitively intact elderly individuals with a Geriatric depression scale (GDS) ^57^ score ≤10, Mini mental status examination (MMSE) ^58^ score ≥25, no current neurological and psychiatric illness, normal functions on verbal and visual memory tests (all scores ≥−1.5 SD of age-adjusted, gender-adjusted, and education-adjusted norms) and age of 60–90 (inclusive) years. All subjects underwent a detailed standardised neuropsychological test session and neuroimaging measurements, all of which were obtained in close temporal proximity with follow up every 1 to 2 years.

Data from 488 individuals from ADNI 2/GO were used to train the machine learning model. Individuals were placed into three categories based on their baseline and longitudinal clinical diagnosis, with baseline defined as the evaluation closest to the first florbetapir (FBP) PET scan acquired in ADNI: **Demented** (n=181, 158 amyloid positive at baseline): individuals have a stable diagnosis of demented; **Stable Condition (SC)** (n=145, 34 amyloid positive at baseline): individuals have a baseline diagnosis of cognitively normal and retain this diagnosis at follow up for 4 or more years (mean=5.7+-std=1 years); **Early Alzheimer’s Disease (EAD)** (n=162, 135 amyloid positive at baseline): individuals have a baseline diagnosis (at date of FBP scan) of either cognitively normal (n=18) or MCI (n=144) but received a diagnosis of demented in future clinical evaluation (i.e. progressed to dementia (n=81), or had been diagnosed as demented in a clinical evaluation prior to baseline (i.e. reverted (n=81). We included individuals in the EAD group who were MCI at baseline but have received a diagnosis of demented prior to baseline (i.e. reverted) in this group as we anticipate they are likely affected by AD pathology but are at an earlier stage of AD than the demented (i.e. late AD) group. Further, as our machine learning model is designed with limited degrees freedom, when training using noisy diagnostic labels it is optimised to account for target uncertainty without leading to over-fitting. Therefore, including the additional training samples (i.e. 81 individuals who reverted to MCI) will likely improve model training even though their diagnostic labels may have poor reliability.

Data from 115 individuals from ADNI 3 were used to test the relationship between the prognostic index and regional future tau accumulation. These individuals were either cognitively normal (n=72) or MCI (n=43) at baseline (defined as the diagnosis closest to the first flortaucipir (FTP) PET scan acquired in ADNI 3) and have at least one follow-up FTP PET scan.

Data from 56 community dwelling individuals from BACS were used to test the accuracy of predictions of regional future tau accumulation. These individuals were cognitively normal (n=56) at baseline (defined as the diagnosis closest to the first FTP PET scan acquired in BACS) and have at least one follow-up FTP PET scan.

### MRI Acquisition

Structural MRIs for the ADNI samples were acquired at ADNI-GO, ADNI-2 and ADNI-3 sites equipped with 3 T MRI scanners using a 3D MP-RAGE or IR-SPGR T1-weighted sequences, as described online (http://adni.loni.usc.edu/methods/documents/mri-protocols). Structural MRIs for the BACS sample were collected on either a 1.5T MRI scanner at Lawrence Berkeley National Laboratory (LBNL) or a 3T MRI scanner at UC Berkeley using 3D MP-RAGE T1- weighted sequences. All ADNI and BACS scans were acquired with voxel sizes of approximately 1mm X 1mm X 1mm. MRI data were used for quantitation of the PET data.

### PET Acquisition

PET imaging was performed at each ADNI site according to standardised protocols. The FBP-PET protocol entailed the injection of 10 mCi with acquisition of 20 min of emission data at 50- 70 min post injection. The FTP-PET protocol entailed the injection of 10 mCi of tracer followed by acquisition of 30 min of emission data from 75-105 min post injection.

For the BACS, PIB PET scans were collected at LBNL. After ∼15 mCi tracer injection into an antecubital vein, dynamic acquisition frames were obtained in 3D acquisition mode over a 90 min measurement interval (4 × 15 s frames, 8 × 30 s frames, 9 × 60 s frames, 2 × 180 s frames, 8 × 300 s frames, and 3 × 600 s frames) after x-ray CT. FTP PET scans were collected following an injection of ∼10 mCi of tracer in a protocol identical to that used for ADNI. All BACS participants were studied on a Siemens Biograph PET/CT.

### Imaging Analysis-MRI: Medial Temporal Grey Matter Density

Structural scans were segmented into grey matter, white matter and Cerebrospinal Fluid (CSF). The DARTEL toolbox ^59^ was then used to generate a study specific template to which all scans were normalised. Following this, individual grey matter segmentation volumes were normalised to MNI space without modulation. The unmodulated values for each voxel represent grey matter density at the voxel location. All images were then smoothed using a 3mm3 isotropic kernel and resliced to MNI resolution 1.5×1.5×1.5 mm voxel size. All structural MRI pre-processing was performed using Statistical Parametric Mapping 12 (http://www.fil.ion.ucl.ac.uk/spm/).

To generate a single index of medial temporal grey matter density we used a voxel weights matrix that was previously derived to generate an interpretable and interoperable disease-specific biomarker ^19^. In brief, a feature generation methodology (partial least squares regression with recursive feature elimination (PLSr-RFE)) was used to apply a decomposition on a set of predictors (T1-weighted MRI voxels) to create orthogonal latent variables that show the maximum covariance with the response variable (memory score). Further, we performed recursive feature elimination by iteratively removing predictors (voxels) that have weak predictive value. The PLSr- RFE procedure results in a voxel weights matrix that is used to calculate a single score of AD related medial temporal atrophy. This index of medial temporal grey matter density has been shown to predict memory deficits, relate to individual tau burden and discriminates stable MCI and progressive MCI individuals ^19^. To generate an individual’s score of medial temporal grey matter density we performed a matrix multiplication of the previously derived voxel weights matrix and each subject’s pre-processed T1 weighted MRI scans.

### Imaging Analysis (ADNI)-PET: FBP (Florbetapir PET) Aβ

FBP data were realigned, and the mean of all frames was used to co-register FBP data to each participant’s structural MRI. Cortical Standardised Uptake Value Ratios (SUVR)s were generated by averaging FBP retention in a standard group of ROIs defined by FreeSurfer v5.3 (lateral and medial frontal, anterior and posterior cingulate, lateral parietal, and lateral temporal cortical grey matter) and dividing by the average uptake from a composite reference region (including the whole cerebellum, pons/brainstem, and eroded subcortical white matter regions) to create an index of global cortical FBP burden (Aβ) for each subject ^60^.

### Imaging Analysis (BACS)-PET: PiB (Pittsburgh Compound B) Aβ

Distribution volume ratios (DVRs) were generated with Logan graphical analysis on the aligned PIB frames using the native-space grey matter cerebellum as a reference region, fitting 35–90 min after injection.

For each subject, a global cortical PIB index was derived from the native-space DVR image coregistered to the MRI using FreeSurfer (5.3) parcellations using the Desikan-Killiany atlas ^61^ to define frontal (cortical regions anterior to the precentral sulcus), temporal (middle and superior temporal regions), parietal (supramarginal gyrus, inferior/superior parietal lobules, and precuneus), and anterior/posterior cingulate regions- ROIs combined as a weighted average. There was no partial volume correction performed.

### Image analysis-PET: FTP (Flortaucipir PET) tau

FTP data were realigned and the mean of all frames used to coregister FTP to each participant’s MRI acquired closest to the time of the FTP-PET. FTP SUVR images were generated by dividing voxel wise FTP uptake values by the average value within a mask of eroded subcortical white matter regions ^33^. MR images were segmented and parcellated into 72 ROIs taken from the Desikan-Killany atlas using Freesurfer (V5.3). These ROIs were then used to extract regional SUVR data from the normalised FTP-PET images. Left and right hemisphere ROIs were averaged to generate 36 ROIs for further analysis. We calculated the future annualised rate of tau accumulation for each of the 36 ROIs either by taking the difference between the follow-up and baseline FTP-PET scans divided by the time interval in years from baseline (when only 2 FTP scans were taken), or fitting a linear least squares fit to 3 or more FTP-PET scans and extracting the parameter estimate for the slope of the ROI SUVR vs. time in years from baseline (when 3 or more FTP scans were taken). In the ADNI 3 sample the average time between FTP-PET scans is 1.22 +- std: 0.38 years with the number of follow-up FTP-PET scans n (2 FTP-PET scans) =93, n (3 FTP-PET scans) =17, n (4 FTP-PET scans) =5. In the BACS cohort the average time between FTP-PET scans is 1.8 +- std:0.65 years with the number of follow-up FTP-PET scans n (2 FTP-PET scans) =37, n (3 FTP-PET scans) =19.

### Predictors and outcomes

Three baseline biological markers related to AD were used as predictors to generate the scalar projection from the machine learning model: a) Cortical amyloid burden (Aβ) measured using either FBP (ADNI) or PiB (BACS) PET, b) medial temporal grey matter density derived from the T1 weighted structural MRI and c) APOE 4 genotype. Previously, we have shown when trained on these baseline data our machine learning approach can predict future changes in cognition for individuals diagnosed as MCI ^19^. Here, we use the same baseline predictors to generate predictions in a new sample of early AD individuals (i.e. individuals who are cognitively normal or MCI at baseline). The primary outcome measure for the predictive models is regional future annualised rate of tau accumulation (SUVR/year). A secondary outcome measure is changes in future cognition over the same time scale as the longitudinal FTP scans, as measured by the Preclinical Alzheimer’s Cognitive Composite (PACC).

To test changes in future cognition for individuals from ADNI 3 we used the previously derived ADNI-PACC measure (adni.loni.usc.edu) ^24^. Of the 115 ADNI 3 individuals with multiple FTP-PET scans 102 individuals had multiple measures of the PACC over a similar time period (within 6 months of the baseline FTP-PET and the final FTP-PET scan). Future annualised change in PACC is calculated by either taking the difference between the follow-up and baseline PACC scores divided by the time interval in years from baseline (when only 2 PACC scores are available), or fitting a linear least squares fit to 3 or more PACC scores and extracting the parameter estimate for the slope of the PACC vs time in years from baseline (when 3 or more PACC scores are available). The average time between PACC testing sessions scans is 1.04 +- std: 0.44 years with the number of follow-up PACC testing sessions n (2 PACC sessions) =82, n (3 PACC sessions) =18, n (4 PACC sessions) =2.

### Prediction Models

*Generalised Matrix Learning Vector Quantization **(GMLVQ)-Scalar Projection***: We previously developed a machine learning approach based on the GMLVQ classification framework: GMLVQ-Scalar Projection ^19^. This approach allows us to derive a continuous prognostic metric (i.e. scalar projection) by training a model based on diagnostic labels.

Learning Vector Quantisation (LVQ) are classifiers that operate in a supervised manner to iteratively modify class-specific prototypes to find boundaries of discrete classes. In particular, LVQ classifiers are defined by a set of vectors (prototypes) that represent classes within the input space. These prototypes are updated throughout the training phase, resulting in changes in class boundaries. For each training example, the closest prototype for each class is determined. These prototypes are then updated so that the closest prototype representing the same class as the input example is moved towards the input example and those representing different classes are moved further away. The Generalised Matrix LVQ (GMLVQ) ^62^ extends the LVQ utilising a full metric- tensor for a more robust (with respect to the classification task) distance measure in the input space. To do this, the metric tensor induces feature scaling in its diagonal elements, while accounting for task conditional interactions between pairs of features (co-ordinates of the input space) (**Supplementary methods:** *Generalised matrix learning vector quantisation*). Previously we trained a GMLVQ model with baseline multimodal data (medial temporal grey matter density, Aβ, APOE 4 genotype) and show that the GMLVQ modelling approach classifies MCI patients into subgroups (progressive vs. stable) with high specificity and sensitivity ^19^.

Extending the binary model, we derived a single prognostic distance measure (scalar projection) that separates individuals based on their longitudinal diagnosis. The GMLVQ-scalar projection approach derives a continuous distance metric from the trained GMLVQ classifier. This continuous distance measure (scalar projection) is how far an individual is from the stable prototype along the dimension that best separates individuals who have stable (i.e. SC) vs unstable (i.e. EAD) diagnosis. This allows the model to learn implicitly a continuous prognostic score for an individual that may be predictive of underlying pathophysiological change. Previously we calculated the scalar projection separating individuals who are stable MCI from progressive MCI showing that the continuous value predicts individualised rates of future cognitive decline ^19^. Here, we apply the same framework on a new sample, deriving the scalar projection on SC vs EAD and making individualised predictions of future regional tau accumulation in EAD populations.

*GMLVQ- scalar projection implementation:* From the training sample the model learns the multivariate relationship between Aβ, medial temporal grey matter density and APOE 4 (metric tensor *Λ*) and the location in multidimensional space that best classifies SC vs EAD individuals (prototype locations: *w*_(*SC,EAD*)_). For any new subject with Aβ, medial temporal grey matter density and APOE 4 (sample vector: *x_i_*) the scalar projection can be calculated by a series of simple linear equations.

1. Transform the sample vector *x_i_* and prototypes *w*_(*SC,EAD*)_ into the learnt space via the metric tensor *Λ*. Note as the metric tensor is learnt in the squared Euclidean space we transform using the square root of the metric tensor (i.e. *Λ*^1/2^)

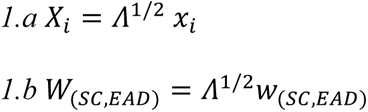

2. Centre the coordinate system on *W*_(*SC*)_ and calculate the orthogonal projection of each vector *X_i_* onto the vector 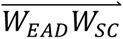, in this co-ordinate system.

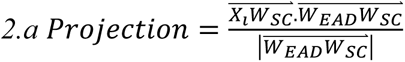

3. To normalise the projections with respect to the position of the prototype *W*_(*SC*)_, the we divided the projection by the norm of 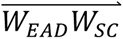:

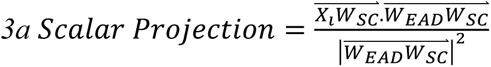

For a graphical derivation and interpretation see **Supplementary Methods**: *GMLVQ-Scalar Projection*.

To determine a meaningful threshold of the scalar projection for separating individuals who are SC from EAD individuals we use logistic regression for the ADNI 2/GO sample labelled as SC, EAD, and Demented. This results in a probabilistic boundary based on the scalar projection.

*Determining Regions of significant AD related tau accumulation:* We first classified individuals from ADNI 3 as either SC or EAD based on each individual’s scalar projection. For the individuals who are classified as EAD (based on the probabilistic threshold value) we performed a subsequent first level analysis to determine which of the 36 selected ROIs will accumulate tau in the future (i.e. regions with a future annualised rate of accumulation statistically greater than 0).

*Predicting individual variability in Regional Tau Accumulation:* Finally, for regions that pass first level significance (i.e. within regions that significantly accumulating tau p<0.05) we trained a series of regression models using ADNI 3 individuals classified as EAD to test if the scalar projection relates to individual variability in regional future rate of tau accumulation (dependent variable: regional future tau accumulation, independent variable: prognostic index). We then tested these models by making individualised predictions –out of sample- for individuals classified as EAD from the BACS sample. To test the accuracy of the regional predictions we calculated the shared variance between the observed future accumulation of tau and the model generated prediction using baseline biological data (i.e. scalar projection).

### Statistical Analysis

Within-sample accuracy for classifying SC vs. EAD in the ADNI2/GO sample was assessed using random resampling. In brief, we determined within sample classification accuracy by randomly splitting our sample into test and training data 400 times. To avoid biasing the model in the training phase due to class imbalance in the data (majority class: EAD= 162 vs. minority class: SC = 145), we resampled the data to generate balanced classes (i.e. number of EAD equals number of SC individuals). This resampling process randomly selects half of the individuals in the minority class and the same number of individuals from the majority class as training data; with the remaining sample used as test data. We then averaged the true positive and true negative accuracies across the 400 resampling’s to generate a class balanced cross validated accuracy.

We used logistic regression to define a probabilistic boundary that separates individuals who are SC from EAD. Using the ADNI 2/GO sample we fit a three class (SC, EAD, Demented) logistic regression to determine the threshold value of the scalar projection. We set the threshold as the probability an individual is less than 50% likely to be SC. To determine the regions that will significantly accumulate tau for individuals classified as EAD we used one tailed one sample t-tests. As we are testing if regions are accumulating tau we use right tail t-tests to determine if the future rate of tau accumulation is significantly (p<0.05) greater than 0 per ROI for individuals classified as EAD. To compare required sample sizes for different models derived using ADNI 3 data we calculated the sample size needed for an arm of a hypothetical clinical trial designed to detect a 25% reduction in annual change (rate of tau accumulation, rate of PACC decline) with a significance of 0.05 and a power of a=0.8. For each comparison, we defined the null hypothesis as the mean and standard deviation of the rate of change calculated from the observed sample, where the alternate hypothesis is a 25% reduction of the mean of the observed sample. For each of the regions that showed significant tau accumulation we fit a robust linear regression (robustfit MATLAB) to predict future tau accumulation using the prognostic index. Setting the dependent variable as future regional tau accumulation and the independent variable as the scalar projection we learnt a series of ROI regression equations.

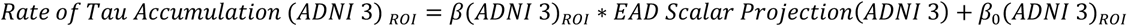

Finally, using the significant (p<0.05) fits derived from ADNI 3 data we generated predictions of tau accumulation for individuals classified as EAD in BACS.

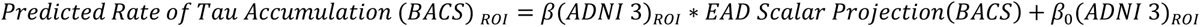

We tested the accuracy of these predictions in the BACS sample by calculating the shared variance between the predicted future rate of tau accumulation and the observed future rate of tau accumulation after treating for outliers (robust correlation ^63^). We determine significance thresholds for the ADNI 3 sample as a p<0.05 uncorrected for multiple comparisons. As we validate our predictions using the BACS sample, we intended to limit type 2 errors by favouring independent validation over conservative multiple comparison correction methods. For regional predictions in BACS we set a significance threshold of shared variance between predicted and observed future rate and tau accumulation as R^2^> 17%. This value corresponds to the coefficient of determination for significance at p<0.05 at the critical value of Pearson’s correlation coefficient (r) with a sample size of 23 (i.e. critical value: r(21)=0.413).

## Supplementary Materials

**Supplementary Table 1. Regional future annualised rate of tau accumulation SC vs EAD**

**Supplementary Table 2. Regional future annualised rate of tau accumulation CN vs MCI**

**Supplementary Figure 1. Regional future annualised rate of tau accumulation CN vs MCI**

**Supplementary Table 3. Regional Future Annualised Rate of Tau Accumulation Early AD vs β-amyloid Positive**

**Supplementary Table 4. Fitting individual variability in regional future annualised rate of tau accumulation**

**Supplementary Table 5. Predicting regional future annualised tau accumulation in BACS**

**Supplementary Methods. Generalised Matrix Learning Vector Quantisation**

## Acknowledgments

We thank Avraam Papadopoulos for help with computational resources. Data collection and sharing for this project was funded by the Alzheimer’s Disease Neuroimaging Initiative (ADNI) (National Institutes of Health Grant U01 AG024904) and DOD ADNI (Department of Defense award number W81XWH-12-2-0012). ADNI is funded by the National Institute on Aging, the National Institute of Biomedical Imaging and Bioengineering, and through generous contributions from the following: AbbVie, Alzheimer’s Association; Alzheimer’s Drug Discovery Foundation; Araclon Biotech; BioClinica, Inc.; Biogen; Bristol-Myers Squibb Company; CereSpir, Inc.; Cogstate; Eisai Inc.; Elan Pharmaceuticals, Inc.; Eli Lilly and Company; EuroImmun; F. Hoffmann-La Roche Ltd and its affiliated company Genentech, Inc.; Fujirebio; GE Healthcare; IXICO Ltd.; Janssen Alzheimer Immunotherapy Research & Development, LLC.; Johnson & Johnson Pharmaceutical Research & Development LLC.; Lumosity; Lundbeck; Merck & Co., Inc.; Meso Scale Diagnostics, LLC.; NeuroRx Research; Neurotrack Technologies; Novartis Pharmaceuticals Corporation; Pfizer Inc.; Piramal Imaging; Servier; Takeda Pharmaceutical Company; and Transition Therapeutics. The Canadian Institutes of Health Research is providing funds to support ADNI clinical sites in Canada. Private sector contributions are facilitated by the Foundation for the National Institutes of Health (www.fnih.org). The grantee organization is the Northern California Institute for Research and Education, and the study is coordinated by the Alzheimer’s Therapeutic Research Institute at the University of Southern California. ADNI data are disseminated by the Laboratory for Neuro Imaging at the University of Southern California.

## Author contributions

JG: Conceptualisation, Formal analysis, Investigation, Methodology, Writing - original draft, Writing - review & editing. WJJ: Conceptualisation, Data curation, Investigation, Writing - original draft, Writing - review & editing. SB: Conceptualisation, Data curation, Formal analysis, Investigation, Writing - original draft, Writing - review & editing .SML: Conceptualisation, Data curation, Formal analysis, Investigation, Writing - original draft, Writing - review & editing. PT: Conceptualisation, Investigation, Methodology, Writing - original draft, Writing - review & editing. ZK: Conceptualisation, Investigation, Methodology, Writing - original draft, Writing - review & editing.

## Data and materials availability

All data and code used in this work are available on request.

---

**Supplementary Table 1.** Regional future annualised rate of tau accumulation. Shows measures of future regional annualised rate of tau accumulation taken from the Desikan Killany atlas for ADNI 3 individuals within the 6 Braak stages. The mean future annualised rate of tau accumulation and test statistics describing whether a region is significantly accumulating tau for individuals classified as Stable Condition (SC) (Left block) and for individuals classified as Early AD (EAD) (Right block). Entries in bold are significant regions for p<0.05 uncorrected.

| Braak Stage | Region | Stable Condition (n=61) |  |  | Early AD (n=54) |  |  |
| --- | --- | --- | --- | --- | --- | --- | --- |
|  |  |  | Significantly Accumulating Tau |  |  | Significantly Accumulating Tau |  |
|  |  | mean accumulation (SUVR/Year) | t-stat | p-val | mean accumulation (SUVR/Year) | t-stat | p-val |
| 1 | ENTORHINAL | 0.0006 | 0.130 | 0.448 | **0.0120** | **2.259** | **0.014** |
| 2 | HIPPOCAMPUS | 0.0009 | 0.233 | 0.408 | -0.0011 | -0.285 | 0.612 |
| 3 | PARAHIPPOCAMPAL | 0.0021 | 0.574 | 0.284 | 0.0049 | 1.149 | 0.128 |
|  | FUSIFORM | 0.0032 | 0.941 | 0.175 | **0.0135** | **2.920** | **0.003** |
|  | LINGUAL | 0.0018 | 0.507 | 0.307 | 0.0033 | 0.899 | 0.186 |
|  | AMYGDALA | -0.0001 | -0.025 | 0.510 | **0.0099** | **2.235** | **0.015** |
| 4 | MIDDLETEMPORAL | **0.0066** | **1.713** | **0.046** | **0.0164** | **3.332** | **<0.001** |
|  | CAUDALANTERIORCINGULATE | 0.0005 | 0.129 | 0.449 | -0.0012 | -0.541 | 0.705 |
|  | ROSTRALANTERIORCINGULATE | 0.0009 | 0.270 | 0.394 | -0.0070 | -2.842 | 0.997 |
|  | POSTERIORCINGULATE | 0.0025 | 0.796 | 0.215 | **0.0056** | **1.741** | **0.044** |
|  | ISTHMUSCINGULATE | 0.0022 | 0.697 | 0.244 | **0.0070** | **2.331** | **0.012** |
|  | INSULA | -0.0003 | -0.103 | 0.541 | 0.0022 | 0.809 | 0.211 |
|  | INFERIORETEMPORAL | 0.0059 | 1.484 | 0.072 | **0.0171** | **3.477** | **<0.001** |
|  | TEMPORALPOLE | -0.0073 | -1.302 | 0.901 | -0.0001 | -0.041 | 0.516 |
| 5 | SUPERIORFRONTAL | 0.0020 | 0.652 | 0.259 | 0.0021 | 0.852 | 0.199 |
|  | LATERALORBITOFRONTAL | 0.0032 | 1.032 | 0.153 | -0.0023 | -0.870 | 0.806 |
|  | MEDIALORBITOFRONTAL | -0.0011 | -0.309 | 0.621 | -0.0041 | -1.439 | 0.922 |
|  | FRONTALPOLE | 0.0038 | 0.818 | 0.208 | 0.0008 | 0.167 | 0.434 |
|  | CAUDALMIDDLEFRONTAL | 0.0033 | 1.096 | 0.139 | **0.0069** | **1.967** | **0.027** |
|  | ROSTRALMIDDLEFRONTAL | 0.0022 | 0.641 | 0.262 | 0.0026 | 0.836 | 0.203 |
|  | PARSOPERCULARIS | 0.0032 | 1.010 | 0.158 | 0.0037 | 1.476 | 0.073 |
|  | PARSORBITALIS | 0.0067 | 1.348 | 0.091 | -0.0021 | -0.418 | 0.661 |
|  | PARSTRIANGULARIS | 0.0035 | 0.920 | 0.181 | 0.0018 | 0.549 | 0.293 |
|  | LATERALOCIPITAL | 0.0045 | 1.007 | 0.159 | **0.0180** | **2.887** | **0.003** |
|  | SUPRAMARGINAL | 0.0000 | 0.010 | 0.496 | **0.0103** | **2.654** | **0.005** |
|  | INFERIORPARIETAL | 0.0024 | 0.721 | 0.237 | **0.0184** | **3.725** | **<0.001** |
|  | SUPERIORETEMPORAL | 0.0016 | 0.464 | 0.322 | 0.0033 | 0.973 | 0.168 |
|  | SUPERIORPARIETAL | 0.0023 | 0.589 | 0.279 | **0.0150** | **3.213** | **0.001** |
|  | PRECUNEUS | 0.0016 | 0.587 | 0.280 | **0.0088** | **2.694** | **0.005** |
|  | BANKSSTS | 0.0039 | 1.200 | 0.117 | **0.0108** | **2.358** | **0.011** |
|  | TRANSVERSETEMPORAL | -0.0042 | -1.259 | 0.893 | -0.0034 | -0.964 | 0.830 |
| 6 | PERICALCARINE | 0.0025 | 0.793 | 0.216 | 0.0021 | 0.549 | 0.293 |
|  | POSTCENTRAL | 0.0013 | 0.400 | 0.345 | 0.0029 | 0.856 | 0.198 |
|  | CUNEUS | 0.0006 | 0.187 | 0.426 | **0.0082** | **1.890** | **0.032** |
|  | PRECENTRAL | 0.0017 | 0.570 | 0.285 | 0.0025 | 0.910 | 0.184 |
|  | PARACENTRAL | 0.0027 | 0.816 | 0.209 | 0.0051 | 1.499 | 0.070 |
|  | Mean | **0.0019** |  |  | **0.0054** |  |  |

**Supplementary Table 2.** Regional future annualised rate of tau accumulation CN vs MCI. Shows measures of future regional annualised rate of tau accumulation taken from the Desikan Killany atlas for ADNI 3 individuals within the 6 Braak stages. The mean future annualised rate of tau accumulation and test statistics describing whether a region is significantly accumulating tau for Cognitively Normal individuals and for MCI individuals (Right block). Entries in bold are significant regions for p<0.05 uncorrected.

| Braak Stage | Region | Cognitively Normal (n=72) |  |  | MCI (n=43) |  |  |
| --- | --- | --- | --- | --- | --- | --- | --- |
|  |  |  | Significantly Accumulating Tau |  |  | Significantly Accumulating Tau |  |
|  |  | mean accumulation (SUVR/Year) | t-stat | p-val | mean accumulation (SUVR/Year) | t-stat | p-val |
| 1 | ENTORHINAL | 0.004 | 0.984 | 0.164 | 0.010 | 1.442 | 0.078 |
| 2 | HIPPOCAMPUS | 0.001 | 0.253 | 0.401 | -0.002 | -0.312 | 0.622 |
| 3 | PARAHIPPOCAMPAL | 0.002 | 0.547 | 0.293 | 0.006 | 1.311 | 0.098 |
|  | FUSIFORM | **0.005** | **1.844** | **0.035** | **0.013** | **2.142** | **0.019** |
|  | LINGUAL | 0.004 | 1.212 | 0.115 | 0.000 | 0.045 | 0.482 |
|  | AMYGDALA | 0.005 | 1.226 | 0.112 | 0.005 | 0.898 | 0.187 |
| 4 | MIDDLETEMPORAL | **0.008** | **2.230** | **0.014** | **0.016** | **2.915** | **0.003** |
|  | CAUDALANTERIORCINGULATE | -0.001 | -0.456 | 0.675 | 0.001 | 0.385 | 0.351 |
|  | ROSTRALANTERIORCINGULATE | -0.004 | -1.745 | 0.957 | -0.001 | -0.152 | 0.560 |
|  | POSTERIORCINGULATE | 0.003 | 0.815 | 0.209 | **0.006** | **2.085** | **0.022** |
|  | ISTHMUSCINGULATE | 0.004 | 1.316 | 0.096 | 0.005 | 1.873 | 0.034 |
|  | INSULA | 0.002 | 0.754 | 0.227 | -0.001 | -0.143 | 0.556 |
|  | INFERIORETEMPORAL | **0.008** | **2.353** | **0.011** | **0.016** | **2.651** | **0.006** |
|  | TEMPORALPOLE | -0.005 | -1.290 | 0.899 | -0.002 | -0.284 | 0.611 |
| 5 | SUPERIORFRONTAL | 0.002 | 0.704 | 0.242 | 0.003 | 0.753 | 0.228 |
|  | LATERALORBITOFRONTAL | 0.002 | 0.773 | 0.221 | -0.002 | -0.412 | 0.659 |
|  | MEDIALORBITOFRONTAL | -0.002 | -0.575 | 0.716 | -0.004 | -1.028 | 0.845 |
|  | FRONTALPOLE | -0.002 | -0.628 | 0.734 | **0.011** | **1.721** | **0.046** |
|  | CAUDALMIDDLEFRONTAL | 0.003 | 1.043 | 0.150 | **0.008** | **2.204** | **0.017** |
|  | ROSTRALMIDDLEFRONTAL | 0.001 | 0.332 | 0.370 | 0.005 | 1.091 | 0.141 |
|  | PARSOPERCULARIS | 0.004 | 1.566 | 0.061 | 0.003 | 0.736 | 0.233 |
|  | PARSORBITALIS | 0.003 | 0.638 | 0.263 | 0.002 | 0.366 | 0.358 |
|  | PARSTRIANGULARIS | 0.003 | 0.942 | 0.175 | 0.002 | 0.520 | 0.303 |
|  | LATERALOCIPITAL | 0.005 | 1.207 | 0.116 | **0.021** | **2.783** | **0.004** |
|  | SUPRAMARGINAL | 0.004 | 1.404 | 0.082 | 0.006 | 1.438 | 0.079 |
|  | INFERIORPARIETAL | **0.007** | **2.034** | **0.023** | **0.015** | **2.645** | **0.006** |
|  | SUPERIORETEMPORAL | 0.002 | 0.509 | 0.306 | 0.004 | 0.966 | 0.170 |
|  | SUPERIORPARIETAL | 0.006 | 1.569 | 0.061 | **0.013** | **2.293** | **0.013** |
|  | PRECUNEUS | **0.006** | **2.177** | **0.016** | 0.003 | 0.897 | 0.188 |
|  | BANKSSTS | **0.007** | **1.858** | **0.034** | **0.008** | **1.828** | **0.037** |
|  | TRANSVERSETEMPORAL | -0.003 | -0.952 | 0.828 | -0.005 | -1.336 | 0.906 |
| 6 | PERICALCARINE | 0.004 | 1.436 | 0.078 | -0.001 | -0.283 | 0.611 |
|  | POSTCENTRAL | 0.001 | 0.388 | 0.350 | 0.004 | 0.862 | 0.197 |
|  | CUNEUS | 0.004 | 1.139 | 0.129 | 0.004 | 1.020 | 0.157 |
|  | PRECENTRAL | 0.001 | 0.295 | 0.385 | 0.004 | 1.227 | 0.113 |
|  | PARACENTRAL | 0.004 | 1.326 | 0.095 | 0.003 | 0.920 | 0.181 |
|  | Mean | 0.003 |  |  | 0.005 |  |  |

**Supplementary Figure 1.**
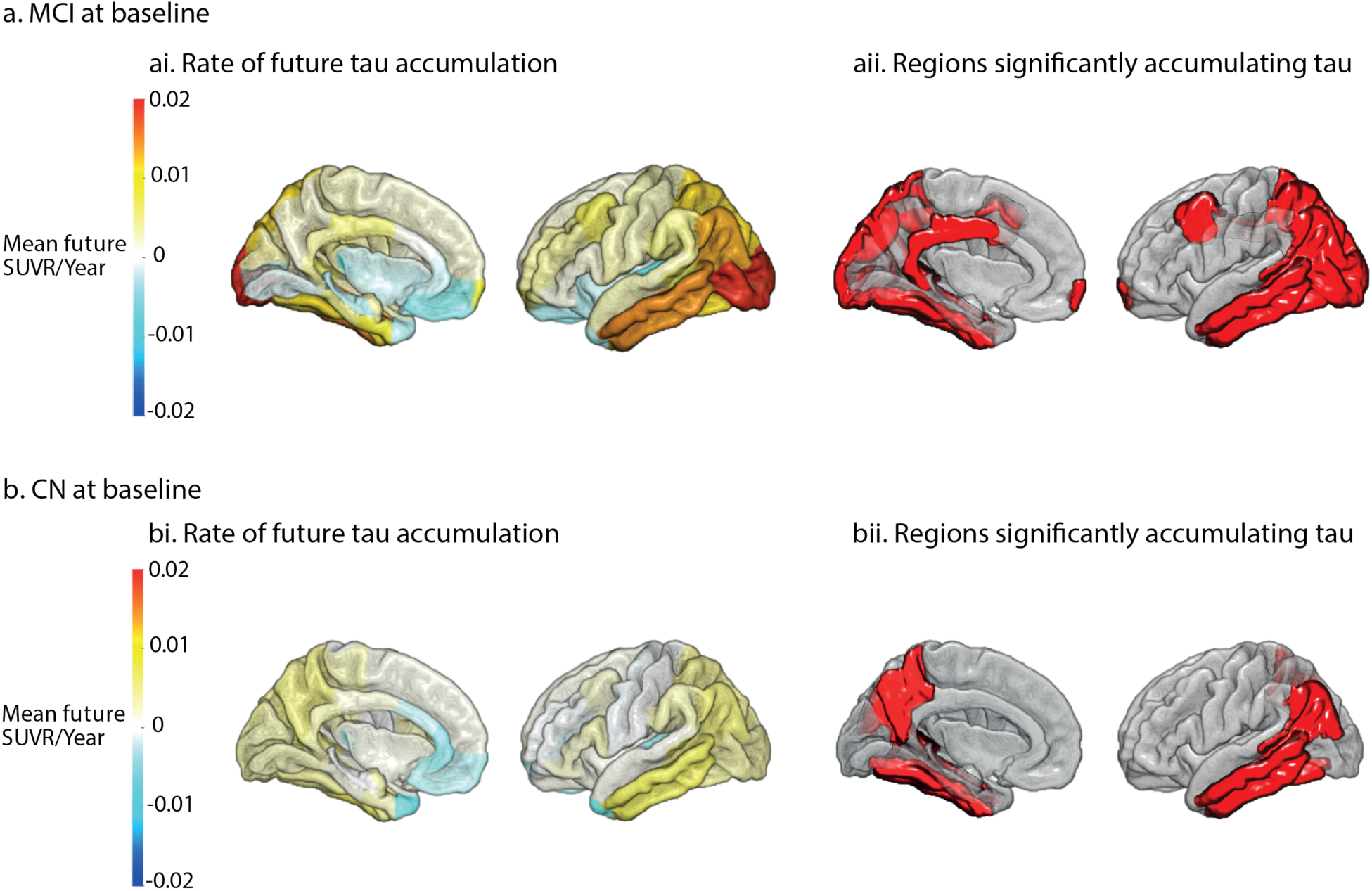
Regional future annualised rate of tau accumulation CN vs MCI. The future annualised rate of tau accumulation for ADNI 3 individuals across the 36 Desikan Kilany ROIs. **ai**. Mean future annualised rate of tau accumulation for MCI. **aii** The regions in red are significantly (p<0.05 uncorrected) accumulating tau for MCI individuals. **bi**. Mean future annualised rate of tau accumulation for Cognitively Normal (CN). **bii** The regions in red are significantly (p<0.05 uncorrected) accumulating tau for Cognitively Normal individuals.

**Supplementary Table 3.**
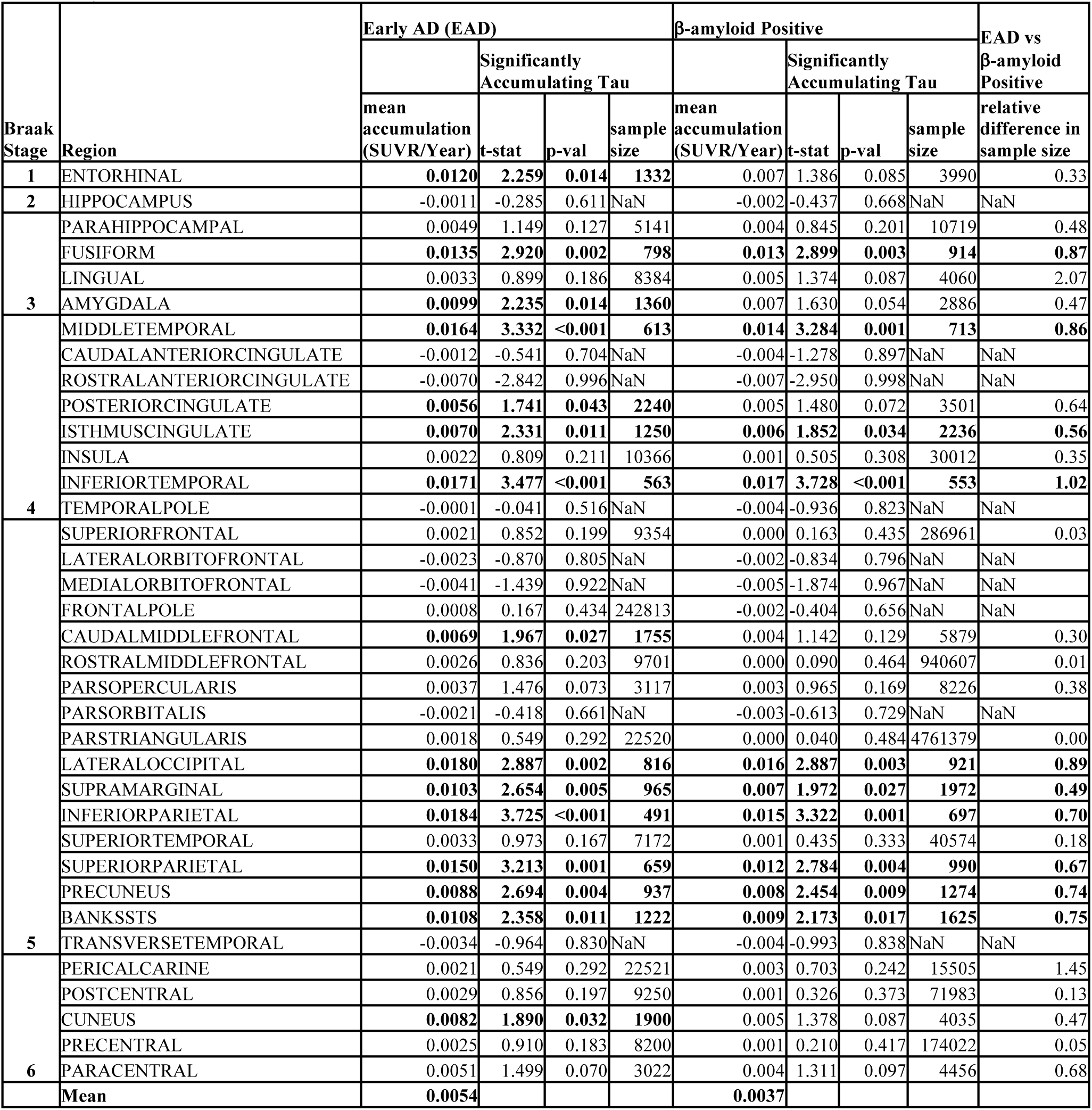
Regional Future Annualised Rate of Tau Accumulation Early AD vs β-amyloid Positive. Shows measures of future annualised rate of tau accumulation for regions taken from the Desikan Killany atlas within the 6 Braak stages. The mean future annualised rate of tau accumulation and test statistics describing whether a region is significantly accumulating tau for individuals classified as Early AD (EAD) (Left block) and for individuals who are β-amyloid positive only (Right block). The Sample size reported is for a hypothetical clinical trial looking to observe a 25% reduction in rate of tau accumulation per region. The right most column compares the required sample size to observe the same effect for groups defined my multimodal data (EAD) or amyloid status only. Entries in bold are significant regions for p<0.05 uncorrected.

**Supplementary Table 4.**
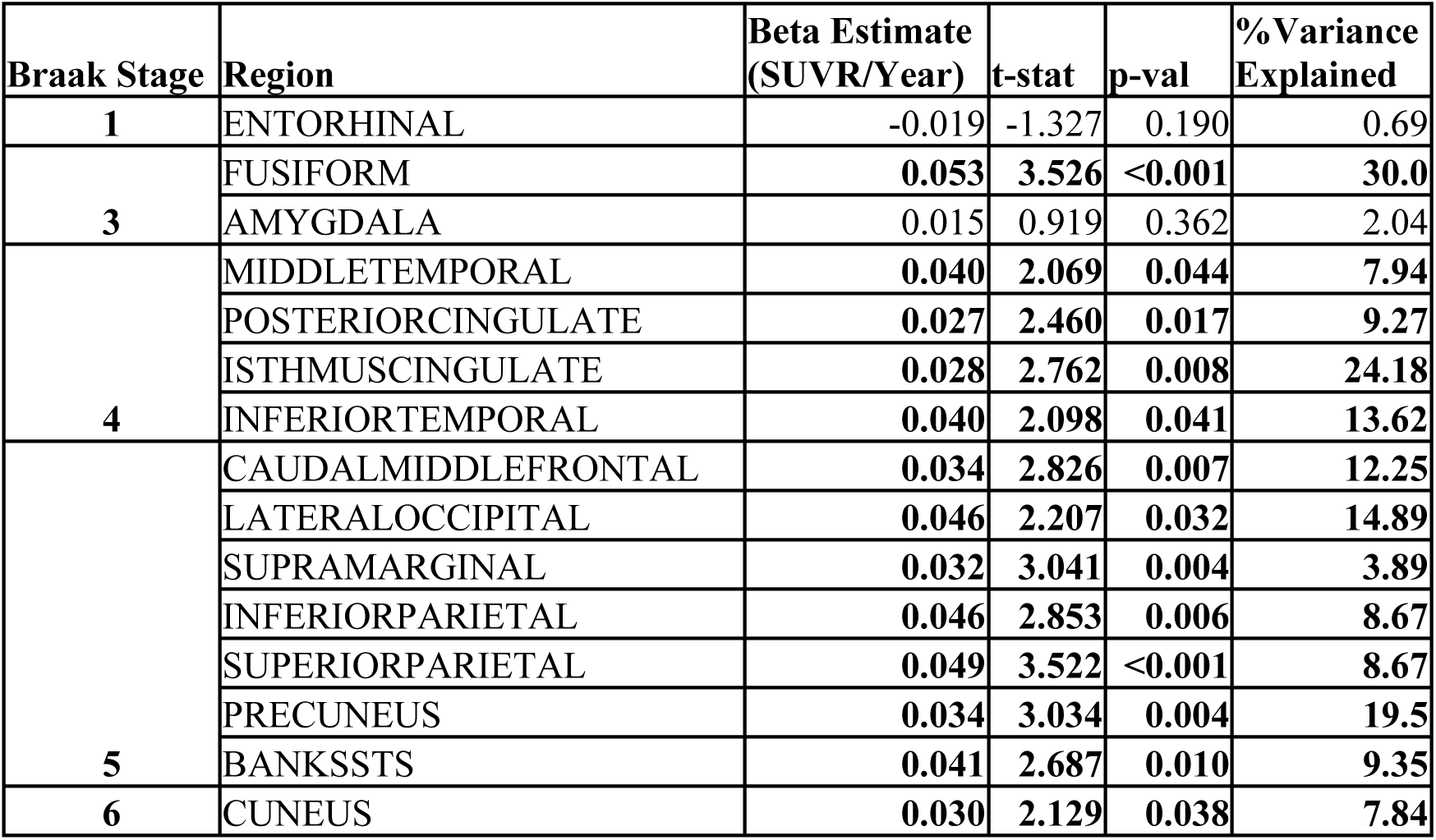
Fitting individual variability in regional future annualised rate of tau accumulation. Shows the parameter estimates and associated statistics for the robust regression equations using the prognostic index to predict regional future tau accumulation for individuals from ADNI 3 classified as Early AD (EAD). Entries in bold are significant regions for p<0.05 uncorrected.

**Supplementary Table 5.** Predicting regional future annualised tau accumulation in BACS. Shows the shared variance of the predicted regional future tau accumulation and the observed future tau accumulation for individuals from the BACS cohort. The right most column shows the shared variance for predicted vs. real tau accumulation for individuals classified as Early AD (EAD). The column second from the right shows the shared variance for predicted vs. real tau accumulation for individuals classified as Stable Condition (SC)

| Braak Stage | Region | ADNI 3 Beta Estimate (SUVR/Year) | SC (n=33) | EAD (n=23) |
| --- | --- | --- | --- | --- |
|  |  |  | Predicted Variance Explained % | Predicted Variance Explained % |
| 3 | FUSIFORM | 0.053 | 3.02 | 26.11 |
| 4 | MIDDLETEMPORAL | 0.040 | 0.48 | 21.82 |
|  | POSTERIORCINGULATE | 0.027 | 0.09 | 12.18 |
|  | ISTHMUSCINGULATE | 0.028 | 0.00 | 2.79 |
|  | INFERIORTEMPORAL | 0.040 | 1.45 | 22.47 |
| 5 | CAUDALMIDDLEFRONTAL | 0.034 | 0.89 | 0.89 |
|  | LATERALOCIPITAL | 0.046 | 2.38 | 6.79 |
|  | SUPRAMARGINAL | 0.032 | 0.28 | 32.19 |
|  | INFERIORPARIETAL | 0.046 | 0.02 | 2.31 |
|  | SUPERIORPARIETAL | 0.049 | 0.59 | 20.95 |
|  | PRECUNEUS | 0.034 | 2.47 | 26.74 |
|  | BANKSSTS | 0.041 | 0.51 | 38.46 |
| 6 | CUNEUS | 0.030 | 2.51 | 0.01 |

